# Annexin A2 and lamin B join membrane recycling compartments for the assembly of biomolecular condensates operating in mitotic partitioning

**DOI:** 10.1101/2024.07.08.602477

**Authors:** Ann Kari Grindheim, Hege Dale, Josef Novák, Patil Sudarshan Shantinath, Anni Vedeler, Jaakko Saraste

## Abstract

Localization of the actin-, lipid- and mRNA-binding protein Annexin A2 (AnxA2) in dividing cells revealed its presence in large spherical structures which are confined to the cell periphery and frequently co-align with astral microtubules. These structures emerge at prometaphase and disappear at telophase, coinciding with the mitotic breakdown and reformation of the nuclear envelope, respectively. Their size increases towards anaphase, while their number decreases, indicating that they are capable of fusion. Treatment of cells with propylene glycol caused rapid and reversible disassembly of the structures, providing additional evidence that they correspond to biomolecular condensates. Interestingly, the condensates contain compartments operating in biosynthetic or endocytic membrane recycling – defined by Rab1, Rab11 or endocytosed transferrin – but are devoid of other membrane organelles, implying that they constitute a mitotic reservoir for selected endomembranes. They also contain lamin B, which connects with the pericentrosomal membrane recycling compartments at prometaphase, when the nuclear lamina breaks down in parallel with centrosome separation. These results suggest that the peripheral condensates correspond to the enigmatic membranous spindle matrix implicated in spindle organization and organelle inheritance. The separating daughter cells at late anaphase contain equal numbers of the condensates, in accordance with their role in mitotic partitioning of endomembranes and other cytoplasmic components.

## Introduction

Cell division involves an extensive reorganization of the cell’s internal architecture and a major change in cell shape (Champion, Linder, and Kutay 2017; Carlton, Jones, and Eggert 2020). Regarding the cytoskeleton, as cells enter mitosis the radial array of microtubules (MTs) typical for interphase cells is reorganized into the mitotic spindle, a bipolar MT-based structure which provides the framework for the alignment and segregation of chromosomes. Astral MTs connecting the spindle poles with the cell cortex, together with the motor protein dynein, determine the correct positioning of the spindle, thereby affecting the fidelity of chromosome segregation (di Pietro, Echard, and Morin 2016). In addition, the cortical network of actin filaments and associated proteins is subject to considerable mitotic remodelling, generating special attachment sites for the astral MTs and facilitating the rounding of the cells (Théry and Bornens 2008; Champion, Linder, and Kutay 2017).

Besides chromosomes also cytoplasmic components, including different organelles of the endomembrane system, must be equally distributed between the daughter cells. However, while the segregation of genetic material is well described, the mechanisms of organelle inheritance remain controversial (Carlton, Jones, and Eggert 2020). Also, how the two processes are coordinated is poorly understood. For correct partitioning, the single copy organelles of the secretory pathway, the endoplasmic reticulum (ER) and Golgi apparatus must undergo remodelling or complete disassembly (Champion, Linder, and Kutay 2017; Ayala, Mascanzoni, and Colanzi 2020). The mitotic ER network is excluded from the region of the spindle (Ellenberg et al. 1997; Schlaitz et al. 2013) and adopts a predominantly tubular or sheet-like organization depending on the cell type (Puhka et al. 2007; Puhka et al. 2012; Lu, Ladinsky, and Kirchhausen 2009; Kumar et al. 2021). A subdomain of the ER – the nuclear envelope (NE) – breaks down at prometaphase jointly with the disassembly of the nuclear lamina. The solubilized A-type lamins are dispersed throughout the cytoplasm, while lipid-linked B-type lamins remain membrane-bound and – like integral NE components – are thought to redistribute to the mitotic ER (Gerace, Blum, and Blobel 1978; Mall et al. 2012; Ungricht and Kutay 2017). However, the release of lamin B, together with that of other nuclear proteins, has also been linked to the formation of a membranous matrix which regulates the proper assembly and function of the mitotic spindle (Tsai et al. 2006; Ma et al. 2009; Zheng 2010; Johansen et al. 2011; Schweizer, Weiss, and Maiato 2014). Although the nature of such a spindle matrix remains enigmatic, its formation is expected to be initiated upon NE break-down and involve dynein-dependent transport of lamin B towards the separating spindle poles (Beaudouin et al. 2002; Salina et al. 2002).

As cells enter mitosis, the Golgi ribbon first undergoes fragmentation, followed by vesiculation of the separated cisternal stacks (Ayala, Mascanzoni, and Colanzi 2020). The disassembled Golgi elements may retain their autonomy and act as a template for organelle reassembly during mitotic exit (Seemann et al. 2002). Alternatively, Golgi enzymes could be recycled back to the ER and – like integral NE proteins – partition as ER components due to the establishment of a mitotic block in ER export (Zaal et al. 1999). Interestingly, in contrast to the Golgi, the intermediate compartment (IC) operating in ER-Golgi trafficking maintains many of its compositional and structural properties, as well as its association with the spindle MTs (Marie et al. 2012). In addition, the IC maintains its connection with the recycling endosomes (REs) at the spindle poles (Hehnly and Doxsey 2014; Marie et al. 2009; Marie et al. 2012; Takatsu et al. 2013), opening for joint partitioning of these pericentrosomal compartments at the onset of mitosis, coupled to centrosome separation and formation of the spindle (Marie et al. 2012; Takatsu et al. 2013; Saraste and Prydz 2019).

Correct orientation of the spindle apparatus depends on a conserved protein complex that mediates the cortical anchoring of astral MTs (di Pietro, Echard, and Morin 2016). One of the components of this complex is AnxA2 (Pascal et al. 2022), a multifunctional protein that interacts with actin and negatively charged phospholipids – such as phosphatidylinositol 4,5-bisphosphate [PI(4,5)P_2_] – in a Ca^2+^-dependent manner (Grieve, Moss, and Hayes 2012; Rescher et al. 2004; Hayes, Merrifield, et al. 2004; Bharadwaj et al. 2013; Gerke, Creutz, and Moss 2005). Based on these properties and its concentration under the plasma membrane (PM) in interphase cells, AnxA2 has been implicated in cortical actin remodelling (Grieve, Moss, and Hayes 2012; Hayes, Rescher, et al. 2004). In addition, it plays a role in mRNA localization and translation, as well as secretory and endocytic membrane traffic (Grieve, Moss, and Hayes 2012; Grindheim et al. 2016; Mickleburgh et al. 2005; Vedeler et al. 2012; Strand et al. 2021; Grindheim et al. 2023; Gerke et al. 2024). In the endocytic pathway AnxA2 has been shown to associate with REs and multivesicular bodies (Mayran, Parton, and Gruenberg 2003; Zeuschner, Stoorvogel, and Gerke 2001; Zobiack et al. 2003; Delevoye et al. 2016), and to be further targeted to the luminal exosomes of the latter (Grindheim et al. 2016; Grindheim and Vedeler 2016; Valapala and Vishwanatha 2011).

The function of AnxA2 is required for successful cell division. It localises to the intercellular bridge connecting the forming daughter cells (Skop et al. 2004) and is necessary at an early stage of cytokinesis (Benaud et al. 2015), possibly related to its interaction with PI(4,5)P_2_-enriched membrane domains and/or association with Rab11-positive REs that function in membrane delivery to the intercellular bridge (Fielding et al. 2005; Benaud and Prigent 2016; Wilson et al. 2005). In the present study we have employed special fixation conditions to address the localization of AnxA2 during the earlier phases of cell division and uncover its association with large spherical structures – up to 3 μm in diameter – transiently appearing in the periphery of mitotic cells between prometaphase and telophase. We present evidence that these structures represent biomolecular condensates, organelles with liquid-like properties that form through phase separation and play important roles in subcellular organization (Banani et al. 2017). The accumulation of biosynthetic and endocytic membrane recycling compartments – the IC and REs – in the peripheral condensates suggests their participation in mitotic storage and segregation of selected endomembranes and control of cell surface area (Boucrot and Kirchhausen 2007). Furthermore, based on their content of lamin B, we propose that the condensates correspond to the membranous spindle matrix that has been proposed to function in spindle regulation and coordination of mitotic partitioning events (Zheng 2010; Schweizer, Weiss, and Maiato 2014; Johansen et al. 2011).

## Results

### Localisation of AnxA2 in mitotic cells

To address the detailed localisation of AnxA2 during cell division we used unsynchronized cultures of normal rat kidney (NRK) cells, which display a high mitotic index and have been commonly used in studies of mitosis (Marie et al. 2012; Beaudouin et al. 2002; Salina et al. 2002; Seemann et al. 2002). Employment of the standard immunofluorescence protocol, based on paraformaldehyde (PFA)-fixation and saponin-permeabilization of cells, demonstrated cortical accumulation of AnxA2 in the mitotic cells, similar to that seen at interphase (Grieve, Moss, and Hayes 2012; Grindheim et al. 2016). Interestingly, the protein was also detected in distinct peripheral structures, whose size and appearance, however, varied greatly between different experiments, indicating their poor preservation (data not shown). To promote the visualization of these structures we turned to paraformaldehyde-lysine-periodate (PLP) fixation (McLean and Nakane 1974) which by providing cross-linking was supposed to promote their structural preservation (Brown and Farquhar 1989). Indeed, following PLP fixation, these AnxA2-positive spherical structures were consistently larger and displayed a typically smooth appearance, particularly when the fixation was carried out on ice (**Fig. 1A**; see Materials and methods). Indicating their authenticity, similar structures were observed in other cultured cell types, including baby hamster kidney (BHK21) cells (**Fig. 2G-I**), rat embryonal fibroblasts (REF52) and human retinal epithelial (RPE-1) cells (data not shown). By contrast, these AnxA2-containing mitotic structures were not detected in transformed human HeLa cells, in agreement with a previous report (Pascal et al. 2022). However, as explained below, the presence of Tfn-positive peripheral puncta may indicate that similar structures also exist in these cells (**Supplementary Fig.1**, open arrows), but are only partially preserved under the fixation conditions used.

**Figure 1.**
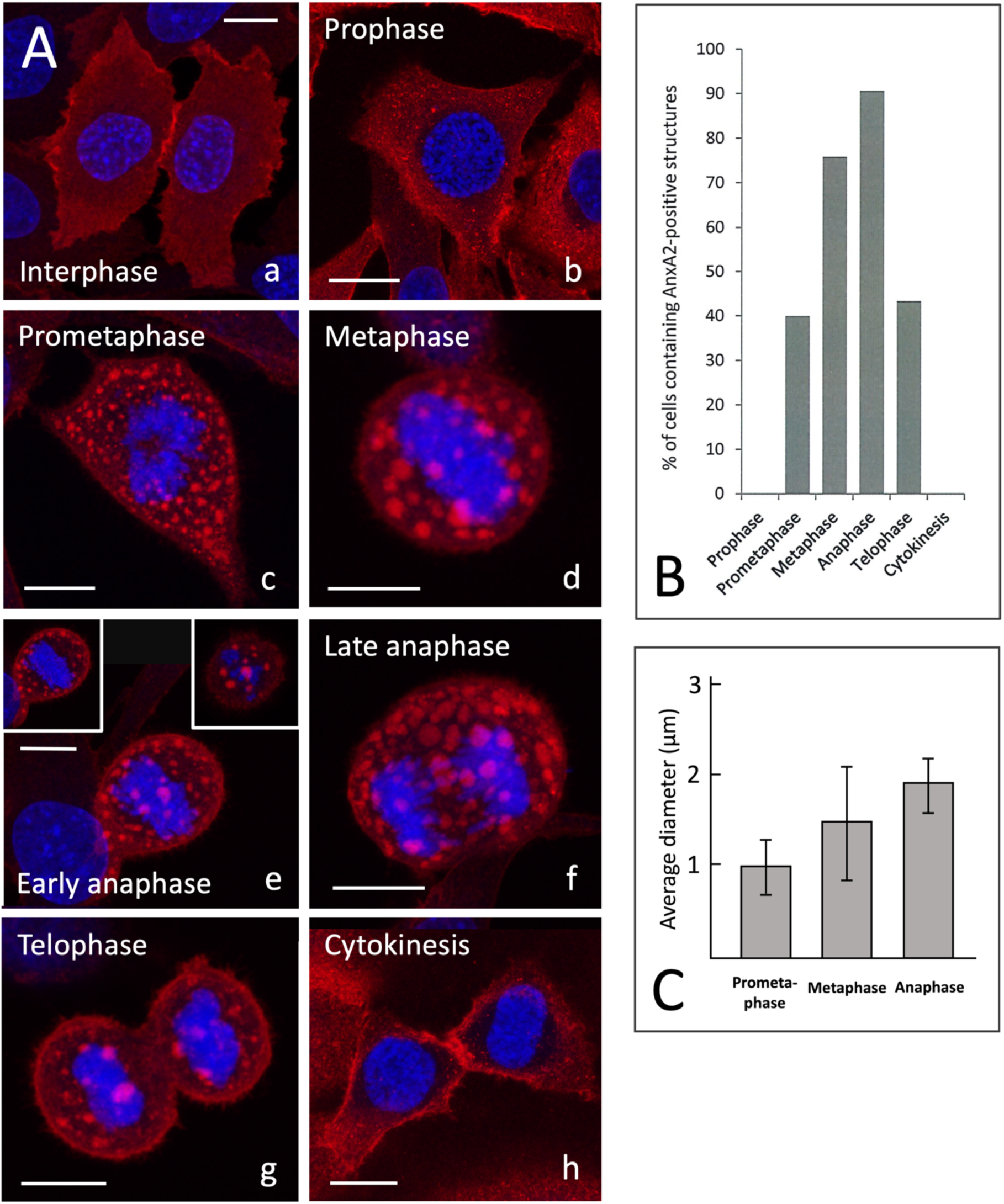
A. Localisation of AnxA2 during the cell cycle. NRK cells were fixed with PLP and permeabilised with saponin. Following the exposure of antigenic sites by treatment with guanidine-HCl the cells were immunolabelled using monoclonal antibodies against AnxA2 (red). Cells at interphase (panel a), the different phases of mitosis (panels b-g), or cytokinesis (panel h) were identified by DAPI staining of DNA (blue). The images represent maximum intensity projections, except the insets in panel e, which show single optical sections from the middle (left inset) or top (right inset) of a cell at early anaphase, demonstrating the characteristic peripheral localization of the large AnxA2-positive structures. Scale bars: 10 µm. B. *Transient appearance of the AnxA2-positive structures during mitosis.* The cells were processed and labelled using monoclonal AnxA2 antibodies and DAPI as described above. The percentages of cells at different phases of mitosis or cytokinesis, positive for the large AnxA2-positive structures, were determined based on the examination of a total of ca. 600 mitotic cells. Note that the great majority of cells at meta- and anaphase are positive for these structures, indicating their structural preservation. C. *The size of the mitotic structures increases during mitosis.* The average size of the structures in prometaphase, metaphase and anaphase cells was determined.

**Figure 2.**
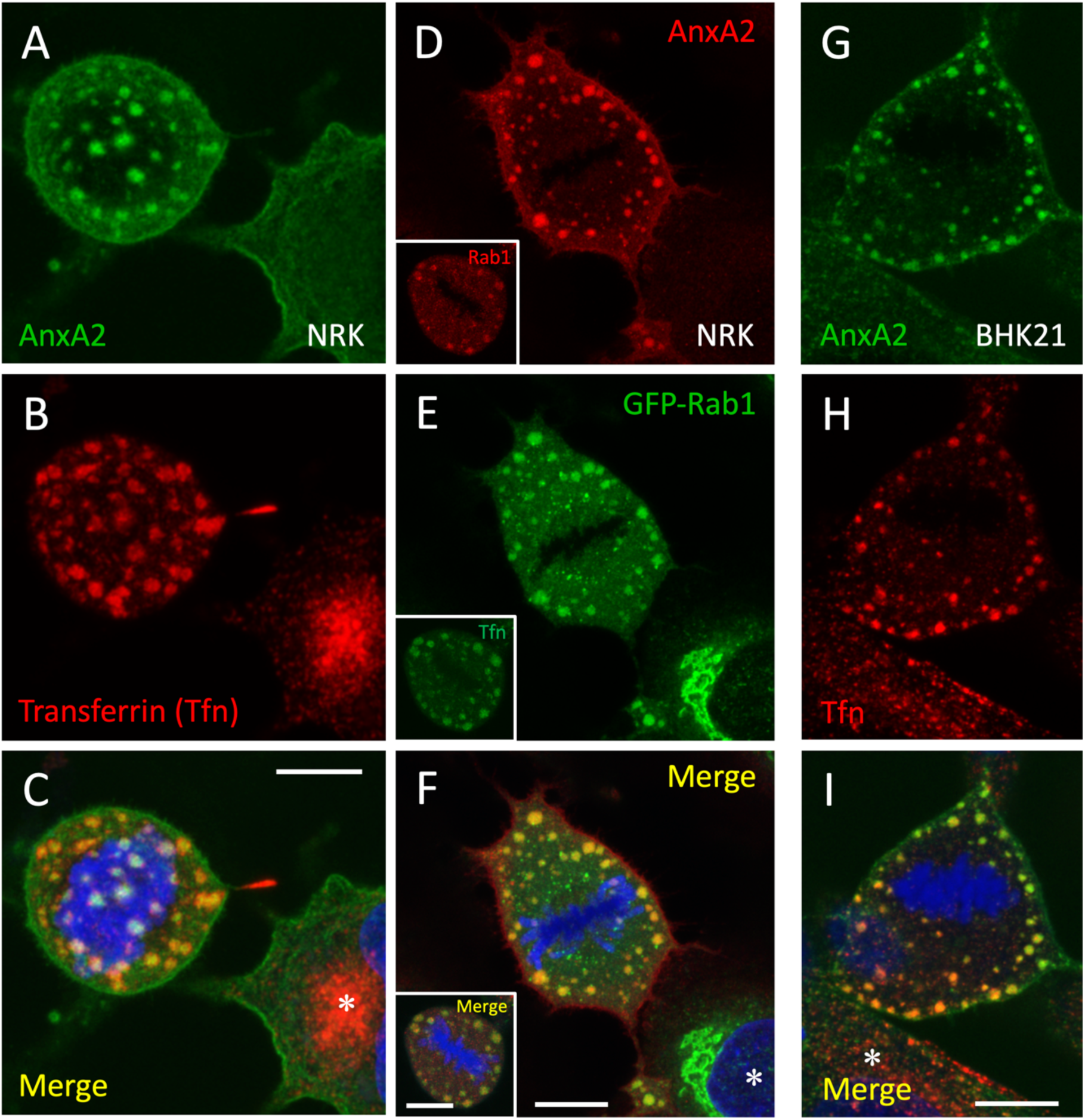
Association of endocytic and biosynthetic membrane recycling compartments with the AnxA2-positive mitotic structures. NRK (panels A-C) or BHK21 cells (panels G-I) were subjected to uptake of Alexa-Fluor 594-transferrin (Tfn; red) to label the recycling endosomal system, followed by fixation with PLP and staining with antibodies against AnxA2 (green) and DAPI (blue) as described for Figure 1. Panels D-F: NRK cells expressing the IC marker GFP-Rab1 (green) were stained for AnxA2 (red). The insets show NRK cells subjected to Alexa-Fluor 488 Tfn-uptake (green) and stained after fixation for endogenous Rab1 (red). The images represent maximum intensity projections from the middle of metaphase cells, while panels A-C show the top of the cell at anaphase. Interphase cells are marked with asterisks. Scale bars: 10 µm.

Examination of the PLP-fixed NRK cells at different phases of the cell cycle confirmed that the novel AnxA2-containing structures are specific for mitotic cells (**Fig. 1A**). To get a better understanding of their nature, we quantified their presence in a large number of dividing cells, employing DNA staining (DAPI) to identify cells at different phases of mitosis or at cytokinesis. As shown in **Fig. 1B**, the structures are absent at prophase, but readily detectable during prometaphase (ca. 40% positive cells). Subsequently, the percentage of positive cells increases from metaphase (ca. 75%) to anaphase (ca. 90%) but decreases again at telophase (ca. 40%). Cells undergoing cytokinesis were devoid of these structures, as also illustrated in **Fig. 1A** (panel **h**). In summary, these AnxA2-positive structures represent transient mitosis-specific assemblies that are detectable between prometaphase and telophase.

The intracellular distribution of AnxA2 in cells at prophase largely resembles that seen during interphase (**Fig. 1A**; compare panels **a** and **b**), while at prometaphase the AnxA2-containing structures – displaying predominantly peripheral localization and variable size – appear concomitantly with the rounding of the cells, (**Fig. 1A**; panel **c**). The generation of Z-stacks of cells at later stages of mitosis demonstrated that these structures are regularly spaced and evenly distributed throughout the cell periphery, mostly lining the cytoplasmic aspect of the PM (**Fig. 1A**; panel **e**, insets). The predominantly peripheral localization of the structures, and their absence from the spindle region is also demonstrated by **Supplementary Movie-1**. Interestingly, their average size increases from 0.9 to 1.95 µm in diameter as the cells progress from prometaphase to anaphase (**Fig. 1C**). The observation that their number concomitantly decreases (see below; **Fig. 6A**) suggests that they are capable of fusion.

### The AnxA2-positive structures contain specific membranes

Based on the well-established association of AnxA2 with the endosomal system (Grieve, Moss, and Hayes 2012; Gerke, Creutz, and Moss 2005; Futter and White 2007), NRK cells were subjected to long-term uptake of fluorescent transferrin which allows the visualization of endocytic membrane compartments at different stages of the cell cycle (Sager, Brown, and Berlin 1984; Tacheva-Grigorova et al. 2013; Takatsu et al. 2013). As previously reported for interphase NRK cells (Marie et al. 2009), internalized transferrin labels both more peripheral endosomal elements, as well as the pericentrosomal REs. Interestingly, the AnxA2-containing structures of the mitotic NRK cells also contained fluorescent transferrin (**Fig. 2A-C**), revealing their connection with the endosomal system. They were also labelled with antibodies against transferrin receptor (data not shown), as well as the GTPase Rab11 (**Supplementary Fig. 4**) – another commonly used marker of REs – indicating that the association of AnxA2 with the endocytic recycling apparatus (Zeuschner, Stoorvogel, and Gerke 2001; Zobiack et al. 2003; Delevoye et al. 2016) is maintained during mitosis.

Furthermore, since the pre-Golgi IC and REs – defined by the GTPases Rab1 and Rab11, respectively – maintain their connection during mitosis (Marie et al. 2012), it was of interest to re-investigate the localization of Rab1 in the mitotic NRK cells fixed with PLP. Interestingly, the employment of cells expressing green fluorescent protein (GFP)-coupled Rab1, or antibodies detecting the endogenous protein, both demonstrated the presence of Rab1 in the mitotic structures containing AnxA2 or transferrin (**Fig. 2D-F**). Of note, while the PLP-fixation helps to visualize endogenous Rab1 in the peripheral mitotic structures, it reduces the strong signal of the protein at the spindle poles (Marie et al. 2012).

We also addressed the presence of other frequently used organelle markers in the novel mitotic structures. As shown in **Supplementary Fig. 2**, regarding organelles of the secretory pathway, the use of antibodies against the ER resident protein calnexin (panels **A-C**), as well as proteins localizing to *cis-*, *medial-* and *trans*-Golgi compartments of the Golgi apparatus – GM130 (panels **D-F**), mannosidase II (panels **G-I**) or TGN46 (panels **J-L**), respectively – failed to show the presence of any of these proteins in the mitotic structures. Moreover, treatment of cells with the drug brefeldin A (BFA), which rapidly releases membrane-bound coat protein I (COPI) coats and results in extensive Golgi disassembly, did not exert any major effect on these structures, suggesting that the Golgi apparatus or the COPI machinery do not play a role in their formation or maintenance (**Supplementary Fig. 3**). Finally, as shown in **Supplementary Fig. 4**, double staining of cells with antibodies against Rab7 (panels **A-C**), EEA1 (panels **G-I**), or LAMP-1 (panels **J-L**) indicated that the structures are also devoid of early or late endosomes, or lysosomes.

Since AnxA2 has been implicated in actin dynamics (Grieve, Moss, and Hayes 2012; Rescher et al. 2008; Hayes et al. 2006; Hayes et al. 2009), it was also possible that the observed mitotic structures could contain aggregates of actin filaments. However, aggregates of comparable appearance were not detected in PLP-fixed mitotic cells stained with the F-actin probe phalloidin. Also, the integrity of the AnxA2-containing structures was maintained in cells treated with the actin filament depolymerizing drug latrunculin B (data not shown). Alter-natively, the structures could be related to the nuclear promyelocytic leukaemia (PML) bodies containing Tyr23 phosphorylated AnxA2 (Grindheim et al. 2016) which during mitosis are released to the cytoplasm and associate with early endosomes marked by the early endosomal antigen 1 (EEA1) (Palibrk et al. 2014). This possibility, however, was excluded by the absence of the PML protein in the AnxA2-positive mitotic structures (data not shown). Finally, the regular spacing of the structures opened the possibility that they associate with the ERM proteins (ezrin, radixin and moesin) which link actin filaments and MTs to the PM (Vilmos et al. 2016). However, the closely PM-associated staining patterns of moesin or pERM antibodies were not reminiscent of the AnxA2-positive structures typically residing at some distance from the PM (data not shown).

In conclusion, these results reveal a specific connection between the AnxA2-containing mitotic structures and compartments operating in membrane recycling at the ER-Golgi boundary (IC) or in the endosomal system (REs). Moreover, they show that in addition to the spindle poles (Marie et al. 2012; Saraste and Prydz 2019) these tubular membrane networks also meet at the periphery of the mitotic cells.

### The mitotic structures co-align with astral MTs

The preferential localization of the mitotic structures to the cell periphery opened the possibility that they are related to the conserved protein complexes that mediate the cortical anchoring of astral MTs and were recently shown to contain AnxA2 (di Pietro, Echard, and Morin 2016; Pascal et al. 2022). Comparison of the distributions of the Rab1-containing large mitotic structures and β-tubulin-containing MTs in cells at anaphase showed that while the former are absent from the central region of the spindle, they frequently co-align with the astral MTs radiating from the spindle poles (**Fig. 3A**, arrowheads). Of note, our previous studies already showed that the Rab1-positive IC elements maintain their association with the MT system during mitosis (Marie et al. 2012).

**Figure 3.**
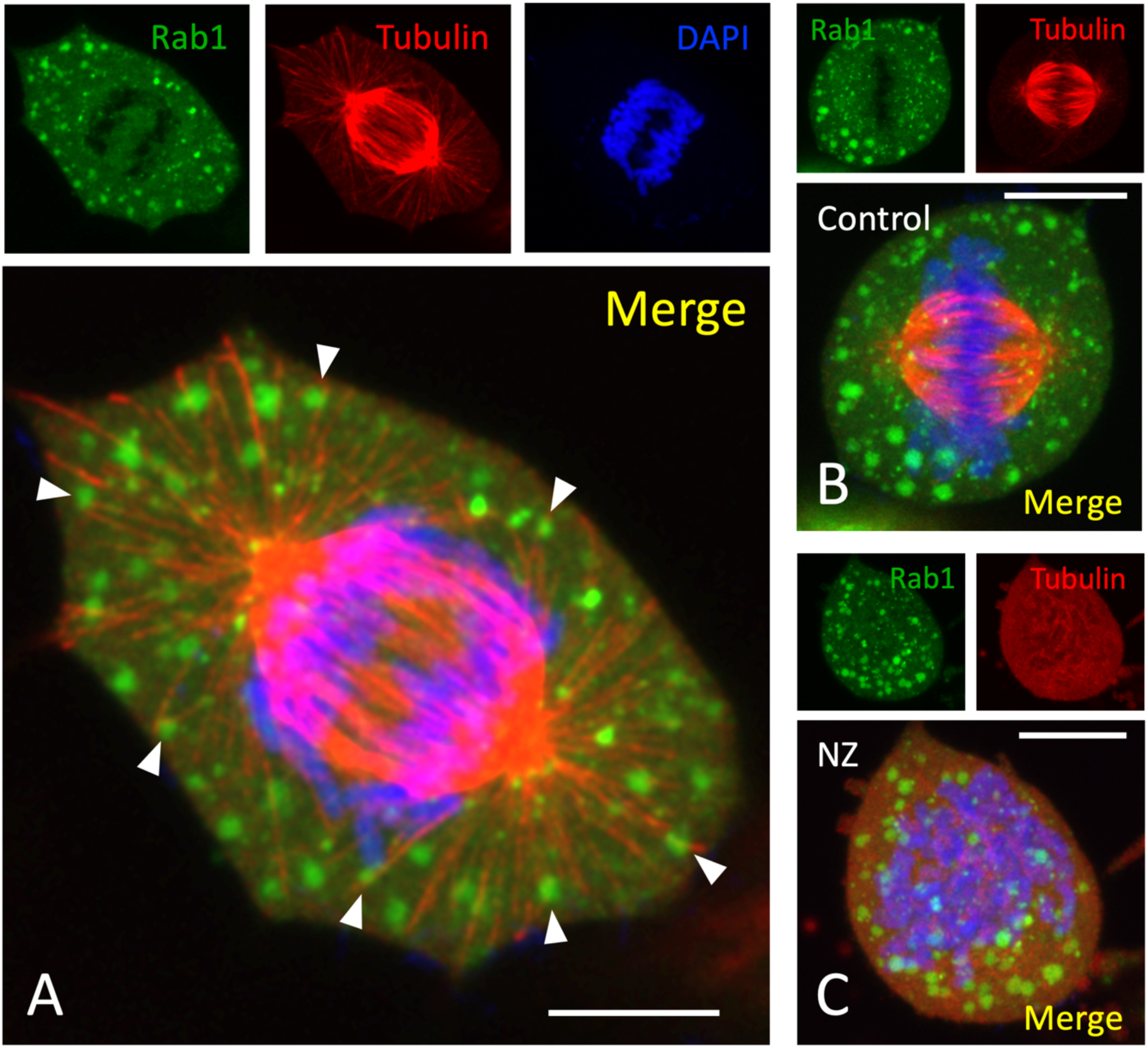
The mitotic structures frequently co-align with astral MTs. NRK cells expressing GFP-Rab1 were fixed with PLP and stained with monoclonal antibodies against β-tubulin and with DAPI. Panel A is an image of a cell at early anaphase, showing frequent co-localization of the large mitotic structures with the astral MTs radiating from the spindle poles (arrowheads) and their absence from the spindle area. Note the accumulation of vesicular Rab1-positive IC elements at the spindle poles. Panels B and C show representative images of similarly stained control metaphase cells (B), or cells treated for 30 min with nocodazole (NZ) to disassemble the spindle MTs (C). Note the dispersal of the large mitotic structures in the drug-treated cells. Scale bars: 10 µm.

To obtain additional information on the functional relationship of the peripheral structures with the mitotic spindle, cells were treated with nocodazole (NZ) to find out whether their structure and/or localization is affected by the disassembly of the spindle MTs. Interestingly, whereas the spherical shape of the structures is unaffected by MT depolymerization, they appear to assume a more dispersed distribution in the drug-treated cells (**Fig. 3B** and **C**). Thus, despite their obvious association with astral MTs, these assemblies are structurally independent of the spindle which, however, may influence their cellular localization.

### The mitotic structures have properties of biomolecular condensates

The large size and spherical shape of the mitotic structures, as well as the need to introduce specific fixation protocols to improve their structural preservation suggested that they may represent biomolecular condensates (Banani et al. 2017), rather than classical membrane-bound organelles. To explore this possibility, we tested their response to aliphatic alcohols, 1,6-hexanediol and 1,2-propanediol – also known as propylene glycol (PG) – which selectively dissolve cellular assemblies formed via liquid-liquid phase separation (LLPS) (Geiger et al. 2021; Kroschwald, Maharana, and Alberti 2017). However, consistent with its known toxicity (Kroschwald, Maharana, and Alberti 2017), 1,6-hexanediol dramatically altered the morphology of mitotic NRK cells, causing many metaphase cells to collapse (data not shown). By contrast, the non-toxic PG is well tolerated by mammalian cells (Geiger et al. 2021; Mochida and Gomyoda 1987) and proved to be applicable also for the investigation of mitotic cells.

Quantitation showed that the addition of PG even at relatively low concentration (2.5%) rapidly reduced the number of metaphase cells containing the mitotic structures and by 5 min they had almost completely disappeared (**Fig. 4A**). Following the removal of PG the percentage of positive cells returned to the initial control level, showing that its cellular effects are readily reversible (**Fig. 4A).** Microscopy of control and PG-treated metaphase cells showed that the breakdown of the AnxA2- and Rab1-containing large structures by PG is not accompanied major changes in cell shape or misalignment of the condensed chromosomes at the equatorial plane (**Fig. 4B** and **C)**. Moreover, PG did not affect the accumulation of the Rab1-positive IC membranes at the spindle poles (Marie et al. 2012); **Fig. 4C**, arrowheads), further indicating that the mitotic spindle is not affected. Strikingly, brief incubation of cells at low temperature gave very similar results to those obtained with PG. Accordingly, in response to 1 to 5 min pre-incubation of cells in ice-cold culture medium, the peripheral mitotic structures gradually disappeared, while the Rab1-positive IC elements at the spindle poles persisted (data not shown).

**Figure 4.**
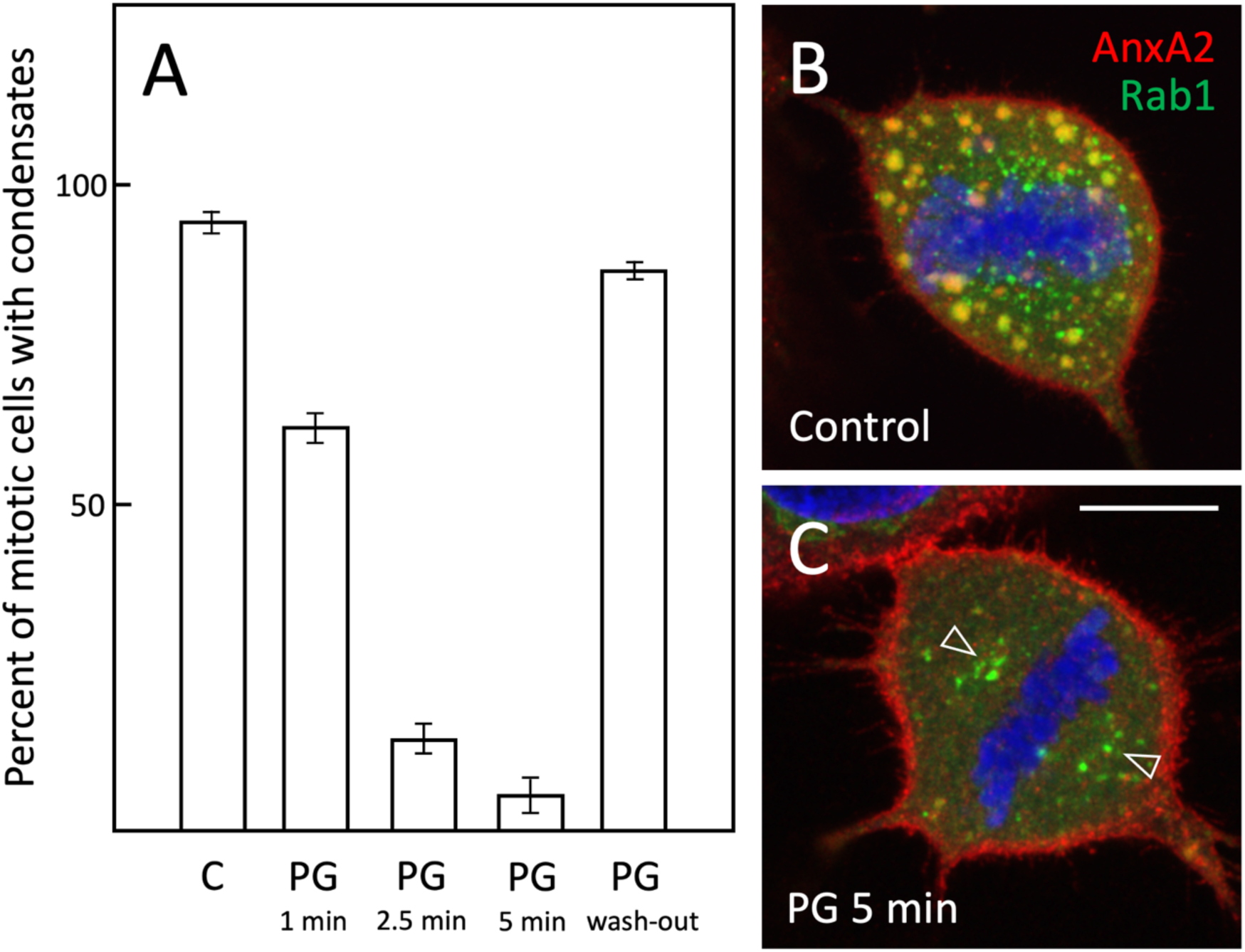
The mitotic structures correspond to biomolecular condensates. A. NRK cells expressing GFP-Rab1 were left untreated (Control), treated for 1–5 min with propylene glycol (PG), or treated for 5 min with PG, followed by a 30 min wash-out of the aliphatic alcohol. After fixation and staining for AnxA2 and DAPI the percentage of cells at meta- or anaphase, containing Rab1- and AnxA2-positive structures were determined. Panels B and C show representative images of control metaphase cells and cells treated for 5 min with PG, respectively. Note the breakdown of the large peripheral structures, increased cortical AnxA2 staining and preservation of the spindle pole-associated IC elements (open arrows) in the PG-treated cells. Scale bar: 10 µm.

In conclusion, the selective breakdown of the large mitotic structures by PG or low temperature strongly suggests that they correspond to biomolecular condensates rather than conventional membrane-bound organelles. However, as presented above, in contrast to the typically membrane-less cytoplasmic biomolecular condensates (Banani et al. 2017) the mitotic condensates identified here contain specific membranes. In this respect they resemble, for instance, the different types of protein condensates found in the presynaptic region of neuronal cells which enclose synaptic vesicles and may closely associate with the PM (Wu, Qiu, and Zhang 2023).

### The mitotic condensates contain lamin B

Biomolecular condensation via LLPS has also been implicated in the assembly of mitotic structures; for example, expansion of the pericentriolar material (PCM) and formation of the spindle matrix, a membranous assembly of nuclear proteins that is independent of MTs, but functionally linked to the mitotic spindle (Woodruff 2018; Tiwary and Zheng 2019). As discussed above, lamin B – one of the protein components that has been employed to define the spindle matrix (Tsai et al. 2006; Ma et al. 2009) – is released as the nuclear lamina breaks down at prometaphase but is thought to maintain its connection with membranes (Champion, Linder, and Kutay 2017). As the novel mitotic structures described in this study evidently represent membrane-associated biomolecular condensates, it was of interest to investigate whether they contain lamin B.

Indeed, double localization of lamin B in metaphase cells with Rab1, transferrin or Rab11 demonstrated its extensive overlap with the markers of the membrane recycling compartments associating with the mitotic condensates (**Fig. 5B** and **C**; **Supplementary Fig. 4**; panels **D**-**F**), providing evidence supporting the idea that they correspond to the membranous spindle matrix defined by lamin B (Zheng 2010). In addition to staining the large peripheral mitotic structures, the lamin B antibodies gave a diffuse cytoplasmic signal which, however, was absent from the spindle area (**Fig. 5C**).

**Figure 5.**
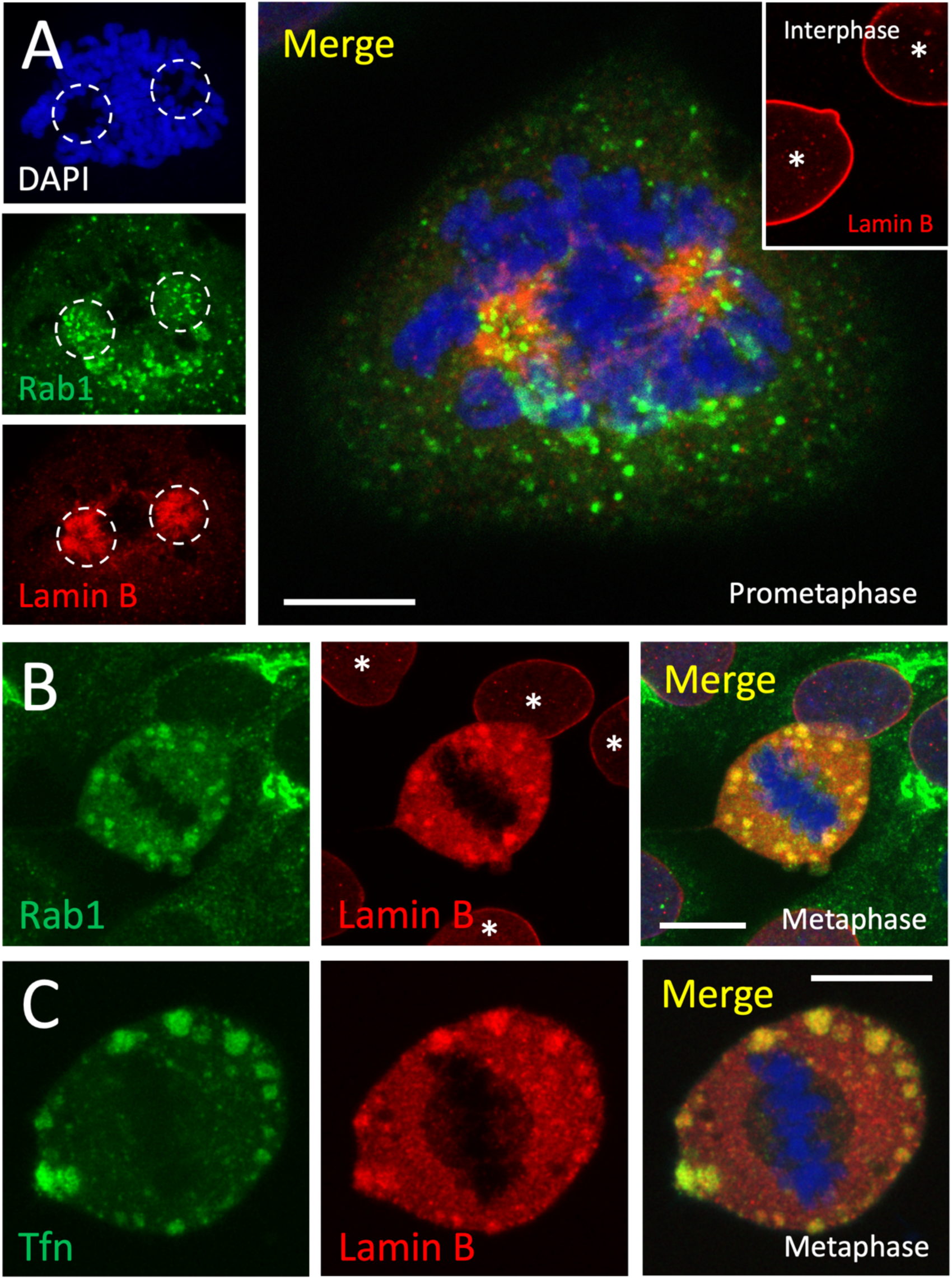
The mitotic structures contain lamin B. NRK cells expressing GFP-Rab1 (panels A and B), or the corresponding parental NRK cells subjected to uptake of Alexa-Fluor 488 Tfn (panel C) were fixed and stained for lamin B (red) and DAPI (blue). Panel A shows a cell at prometaphase, following disassembly of the nuclear lamina, in which both lamin B and the Rab1-positive pericentrosomal IC elements pile up at the separating spindle poles residing in nuclear invaginations revealed by the DAPI-staining (dashed circles). The inset shows lamin B staining of interphase cells (asterisks). Panels B and C demonstrate the co-localization of lamin B with Rab1 and Tfn in the large mitotic structures at the periphery of metaphase cells. Note the additional diffuse cytoplasmic staining for lamin B and its absence – in appropriate projections (panel C) – from the spindle area. Scale bars: 5 µm (A), 10 µm (B and C).

To gain insight into the pathway that lamin B follows to reach the peripheral structures, we examined cells at prometaphase, when the NE and the nuclear lamina break down in a process that is intimately coupled to centrosome separation and formation of the bipolar mitotic spindle (Champion, Linder, and Kutay 2017). Namely, previous studies of NRK cells had shown that at the onset of mitosis both lamin B (Beaudouin et al. 2002) and the pericentrosomal membrane recycling compartments, defined by Rab1 and Rab11 (Marie et al. 2012), congregate around the forming spindle poles. Indeed, at the time when the separating centrosomes localize to deep invaginations of the NE (Beaudouin et al. 2002; Salina et al. 2002; Robbins and Gonatas 1964), both lamin B and the Rab1-positive IC elements were seen to simultaneously accumulate around the spindle poles, displaying overlapping distributions (**Fig. 5A**). This suggests that the transfer of lamin B from the disintegrating nuclear lamina to the peripheral condensates emerging at prometaphase is due to its association with the membrane recycling compartment(s) at the spindle poles.

### Equal mitotic partitioning of the condensates

As discussed earlier, as cells progress through mitosis the size of the condensates increases at the same time as their number appears to decrease. Determination of the average number of condensates in cells at different phases of mitosis – between prometaphase and late anaphase – showed that it drops to about half between prometaphase and metaphase, evidently due to their active fusion, but levels out at anaphase (**Fig. 6A**). Interestingly, at late anaphase, the separating daughter cells clearly delineated by the developing cleavage furrow (**Fig. 6B** and **C**) contain equal amounts of the condensates (**Fig. 6A**; inset), indicating that their partitioning is a tightly controlled process.

**Figure 6.**
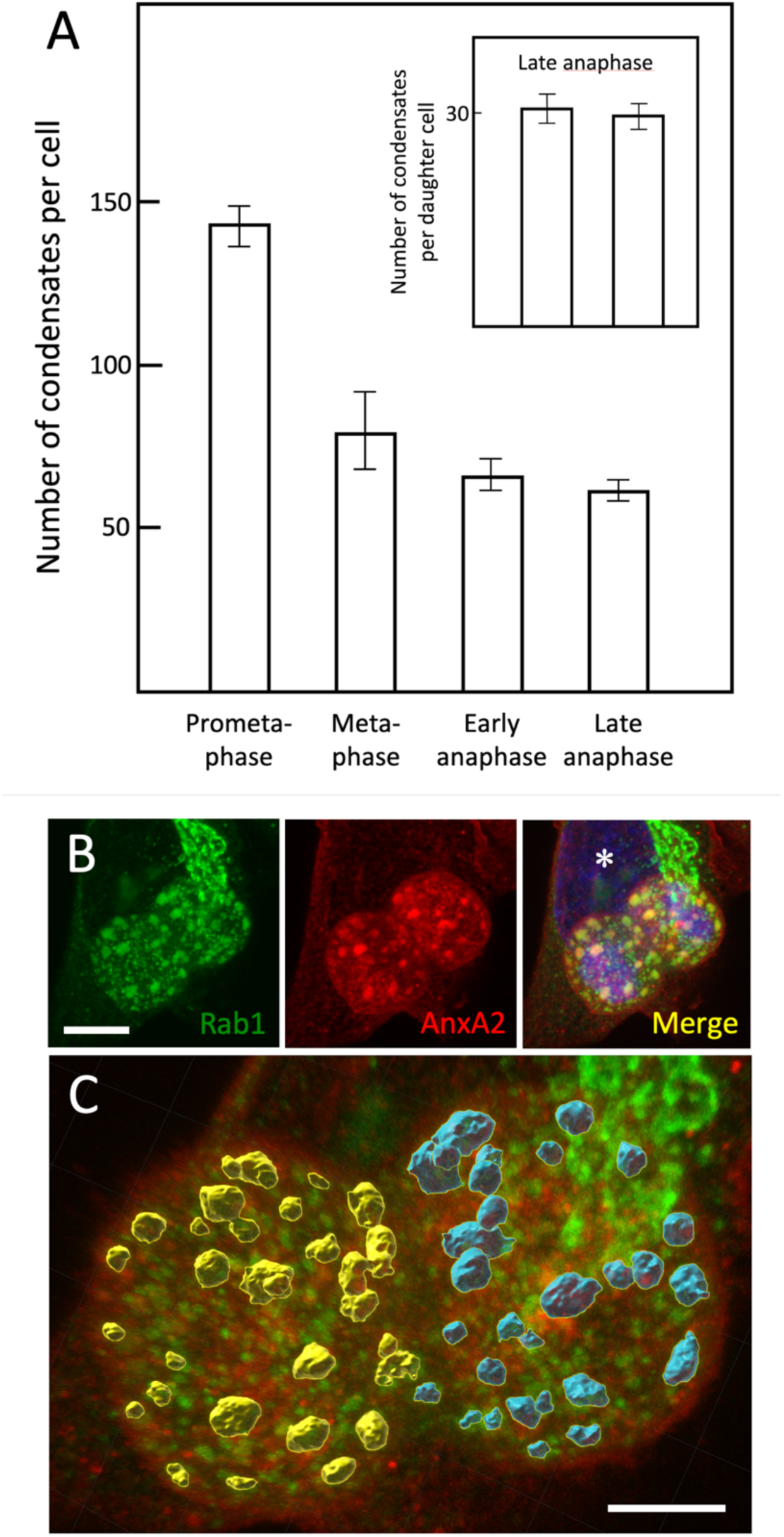
The number of the condensates decreases during mitosis. Panel A: Following fixation and staining of GFP-Rab1-expressing cells for AnxA2, the average number of the condensates in cells at different phases of mitosis was determined. The inset shows similar quantitation carried out for the two separating daughter cells at late anaphase. Panels B and C show representative images of a late anaphase cell, in which the borders of the daughter cells can be readily identified due to the constriction creating by the developing cleavage furrow. An interphase cell is marked by an asterisk. Panel C, corresponding to **Supplementary Movie-2**, was generated by the Imaris software and illustrates the pseudo-coloured large Rab1- and AnxA2-positive condensates in the two daughter cells. Scale bars: 10 µm (B), 5 µm (C).

## Discussion

The present investigation, adopting specific fixation protocols to determine the localization of AnxA2 during cell division, led to the identification of a novel organelle at the periphery of mitotic cells. The exceptionally large size and spherical shape of these structures, as well as their apparent ability to fuse are hallmarks of biomolecular condensates forming by phase separation (Banani et al. 2017). Their rapid and selective breakdown by the aliphatic alcohol propylene glycol (PG), which is expected to influence weak hydrophobic protein-protein and protein-RNA interactions (Geiger et al. 2021), provided additional evidence that they represent biomolecular condensates, rather than classical membrane-bound organelles. In addition, they share similarity with the spindle matrix – a controversial membranous assembly of nuclear proteins, including the nuclear lamina subunit lamin B – whose formation has been suggested to involve phase separation (Schweizer, Weiss, and Maiato 2014; Woodruff 2018; Tiwary and Zheng 2019). Therefore, our results – particularly the localization of lamin B to the membrane-containing mitotic structures – lead us to conclude that they correspond to the previously proposed spindle matrix (Zheng 2010).

A general idea is that a membranous spindle matrix surrounds and mechanically supports the spindle apparatus, controlling its proper assembly, orientation and function (Zheng 2010; Johansen et al. 2011; Schweizer, Weiss, and Maiato 2014). Notably, information regarding the mitotic roles of the presently identified protein components of the condensates also indicates their similarity with the spindle matrix. Earlier studies by Zheng and coworkers already demonstrated the interaction of lamin B with various spindle assembly factors, including NuMA and the MT-based motor proteins dynein and kinesin Eg5 (Tsai et al. 2006; Ma et al. 2009). By interacting with actin and PI(4,5)P_2_ (Harder et al. 1997; Hayes et al. 2009; Rescher et al. 2004; Gerke, Creutz, and Moss 2005) AnxA2 possesses properties expected of a protein involved in the “locking” of astral MTs to regularly spaced foci at the cell periphery (Sandquist, Kita, and Bement 2011). Indeed, it was recently found to collaborate with Ahnak, NuMA and dynein in cortical anchoring of astral MTs, facilitating spindle positioning (Pascal et al. 2022). The GTPase Rab11 has been shown to maintain its association with REs during mitosis (Hehnly and Doxsey 2014; Marie et al. 2012; Hobdy-Henderson et al. 2003) and control dynein-dependent transport of key PCM components, such as γ-tubulin and pericentrin, to the spindle poles, thereby promoting the nucleation of astral MTs and correct orientation of the spindle (Hehnly and Doxsey 2014). Moreover, studies of knock-out mice showed that the Rab11A and Rab11B isoforms redundantly regulate spindle function in dividing epithelial progenitor cells (Joseph et al. 2023). Finally, depletion of the IC-associated Rab1 has been reported to affect centrosome maturation and spindle assembly, besides leading to endomembrane alterations in mitotic cells of the *Drosophila* embryo (Rollins and Blankenship 2023).

The extensive reorganization of endomembranes taking place during mitosis has been thought to be accompanied by inhibition of membrane traffic (Fielding and Royle 2013; Shorter and Warren 2002). However, more recent studies indicate that endocytosis continues throughout mitosis, although the rate of endocytic uptake – possibly via different pathways – appears to slow down from prometaphase to anaphase (Tacheva-Grigorova et al. 2013; Aguet et al. 2016; Hinze and Boucrot 2018). By contrast, it has been proposed that endocytic membrane recycling is temporarily arrested between prophase and late anaphase, providing a simple mechanism for mitotic regulation of cell surface area (Boucrot and Kirchhausen 2007; Devenport et al. 2011). It is likely that the mitotic effects on different steps of membrane traffic are essentially coupled to the development of the condensates into a repository for biosynthetic (IC) and endosomal membranes – connecting the two tubular networks that also meet at the spindle poles (Marie et al. 2012; Saraste and Marie 2018) (see **Fig. 7)**. Upon their disassembly at telophase the condensates could provide the membrane source for expansion of the cell surface at the intercellular bridge, explaining the localization of both AnxA2 and Rab11 at this site (Takahashi et al. 2011; Skop et al. 2004).

**Figure 7.**
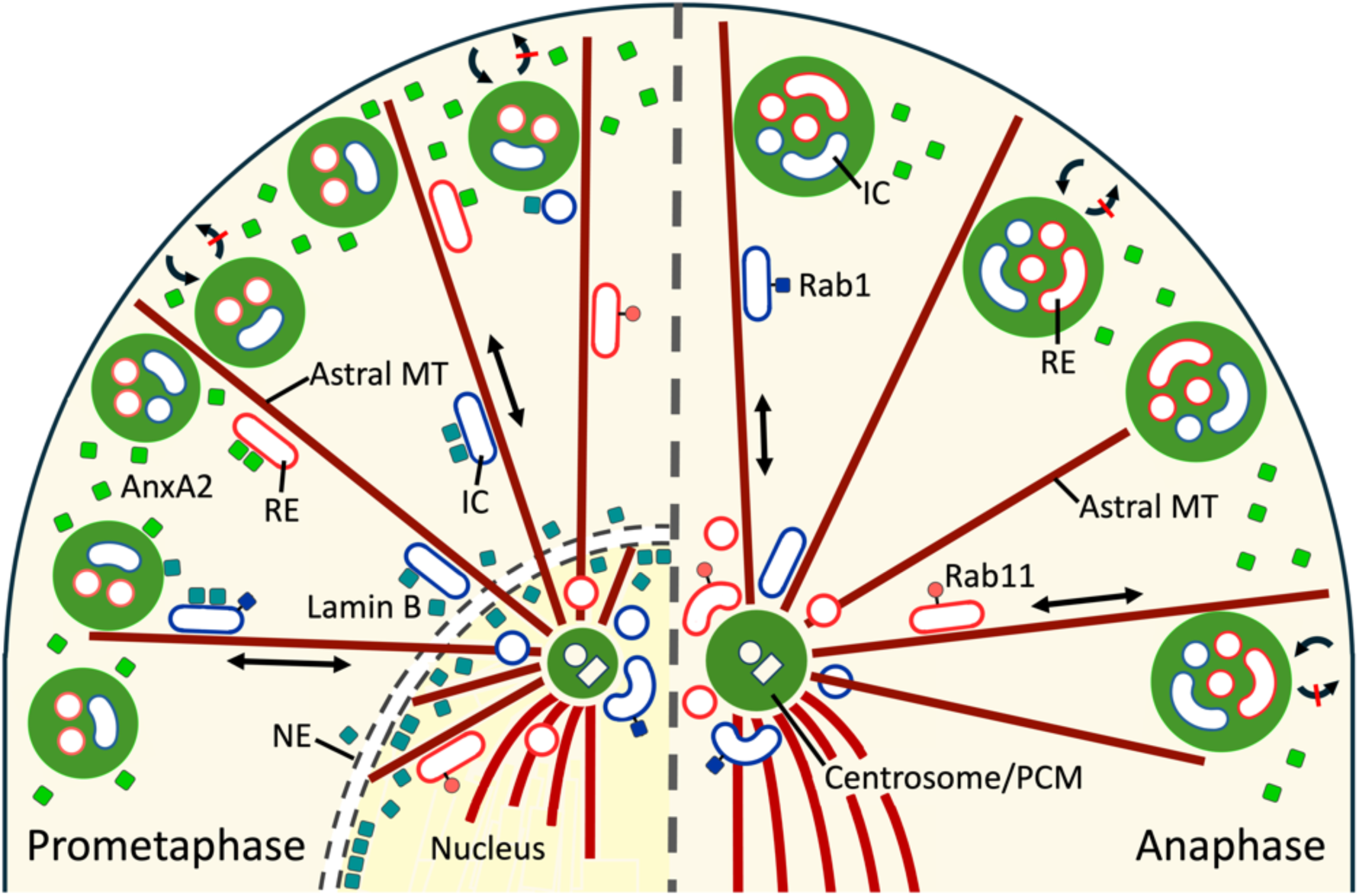
A schematic model on the dynamic events proposed to be implicated in the bio-genesis of the AnxA2-containing mitotic condensates. The illustrated sections of prometa-phase (left) and anaphase cells (right) separated by a dashed line include only one of the spindle poles. At prometaphase the emergence of the AnxA2- and lamin B-containing peripheral condensates, depicted by green spheroids, coincides with the breakdown of the nuclear envelope (NE) and lamina. The released lamin B could move from the spindle poles to the peripheral condensates bound to Rab1-positive IC elements (as shown here) or Rab11-positive REs, based on their motor-dependent trafficking along astral MTs. The recruitment of AnxA2 to the condensates may occur from the cortical cytoplasm or involve its association with the REs. The accumulation of the membrane recycling compartments in the condensates could be due to their proposed communication with the pericentrosomal IC elements and REs at the spindle poles. In addition, the pile-up of REs could result from ongoing endocytic traffic from the cell surface, combined with inhibition of membrane recycling. Two-way communication between the growing condensates and the spindle poles may also be linked to the expansion of the pericentrosomal material (PCM), which has been proposed to involve phase separation.

The discovery of mitotic functions of selected Rab proteins – commonly considered as master regulators of membrane traffic – has revealed that certain transport steps remain largely unaffected during mitosis (Capalbo et al. 2011; Serio et al. 2011; Miserey-Lenkei and Colombo 2016). Together with the above discussed observations regarding Rab11 (Hehnly and Doxsey 2014), the present results open the possibility for ongoing motor-dependent transport along the astral MTs connecting the peripheral condensates with the spindle poles, as suggested by the model shown in **Fig. 7**. Indeed, the presence of the two membrane recycling compartments both at the peripheral condensates and the spindle poles supports the idea that these sites communicate via membrane traffic (**Fig. 7**). Since phase separation has been implicated both in the assembly of the spindle matrix and the expansion of the PCM (Woodruff 2018; Tiwary and Zheng 2019), it is tempting to speculate that the events taking place at the plus and minus ends of astral MTs may involve the exchange of key components (**Fig. 7**). For example, NuMA, which associates with AnxA2 and lamin B, and is found both at the cell cortex and the spindle poles, turns out to regulate spindle assembly and function via phase separation (Ma et al. 2009; Sun et al. 2021). Moreover, lamin B is expected to employ the astral MTs as tracks during its transfer from the spindle poles to the peripheral condensates as the nuclear lamina breaks down at prometaphase (**Fig. 7**).

In interphase cells the local concentration of proteins and nucleic acids caused by bio-molecular condensation is utilized to give rise to organelles with distinct synthetic activities and functions (Banani et al. 2017). Since cellular activities are generally slowed down during mitosis, the concentration of cellular constituents via phase separation could also be utilized for their accurate partitioning. The finding that the forming daughter cells at late anaphase contain equal numbers of the condensates provides evidence that they play a role in this process. For example, the presence of lamin B in these structures suggests that they contribute to the mitotic segregation of nuclear proteins. The role of AnxA2 as an mRNA-binding protein regulating mRNA transport and translation of specific mRNAs (Grieve, Moss, and Hayes 2012; Grindheim et al. 2016; Grindheim, Saraste, and Vedeler 2017; Mickleburgh et al. 2005; Vedeler and Hollas 2000; Vedeler et al. 2012; Strand et al. 2021; Grindheim et al. 2023) opens the possibility that the condensates act in the partitioning of selected mRNAs. Finally, the striking identification of the biosynthetic and endocytic recycling systems as membrane components of mitotic condensates suggests a role of phase separation in the division of endomembranes.

The assembly and disassembly of biomolecular condensates can be regulated by phosphorylation and/or dephosphorylation of their key protein components (Sridharan et al. 2022). Since many of the cellular functions of AnxA2 are regulated by phosphorylation (Grindheim, Saraste, and Vedeler 2017), we assessed the pTyr23 and pSer25 status of AnxA2 in mitotic vs. interphase NRK cells but observed no apparent change in the phosphorylation pattern of the protein (data not shown). Thus, the most likely central role of AnxA2 in the mitotic condensates does not seem to involve its phosphorylation and remains a subject of future work.

## Materials and methods

### Antibodies and reagents

Mouse monoclonal antibodies against AnxA2 (1:200 dilution), early endosomal antigen 1 (14/EEA1; 1:50 dilution), Rab11 (47/Rab11; 1:50 dilution), GM130 (35/GM130; 1:200 dilution) and TGN38 (2/TGN38; 1:50 dilution) were purchased from BD Transduction Laboratories. Rabbit polyclonal antibodies against AnxA2 (ab41803; 1:250 dilution) and Lamin B (ab16048; 1:200 dilution) were obtained from Abcam. The mouse monoclonal transferrin receptor antibody (H68.4; 1:200 dilution) and rabbit polyclonal Rab11 antibody (71-5300; 1:50 dilution) were from Invitrogen, while the mouse monoclonal antibody against LAMP-1 (H5G11; 1:200 dilution) and rabbit polyclonal antibody against Rab7 (R4779; 1:50 dilution) were purchased from Santa Cruz Biotechnology and Sigma, respectively. The mouse monoclonal antibody against Rab1B (1:200 dilution) was generously provided by Angelica Barnekow (University of Münster, Germany), while the rabbit polyclonal antibodies against calnexin (1:100 dilution) and mannosidase II (1:500 dilution) were generous gifts from Ari Helenius (ETH, Zürich, Switzerland) and Kelley Moremen (University of Georgia, USA), respectively. The mouse ascites fluid against β-tubulin (T13; 1:500 dilution) was provided by the late Thomas Kreis. The primary antibodies were detected using appropriate Alexa Fluor 488- or Alexa Fluor 594-conjugated secondary goat anti-mouse or anti-rabbit Fab_2_-fragments (1:100 dilution) bought from Jackson Immuno-Research Laboratories. Alexa Fluor 488- and Alexa Fluor 594-conjugated human transferrin were obtained from Invitrogen. Brefeldin A (BFA), nocodazole (NZ) and propylene glycol (PG) were purchased from Sigma.

### Cell culture

Normal rat kidney (NRK) cells and NRK cells stably expressing the GFP-Rab1 fusion protein (Marie et al. 2009) were grown in Dulbecco’s Minimum Essential Medium (DMEM) supplemented with 10% heat-inactivated fetal calf serum (FCS), 2 mM L-glutamine, 50 units/ml penicillin and 50 μg/ml streptomycin. Mitotic synchronization with drugs was not employed to avoid possible secondary effects. To obtain steady state cultures with high mitotic index, 100% confluent cultures of cells were diluted 1:2 and plated on 18 mm diameter glass coverslips in 6-well plates, followed by growth for 22-24 hr. Baby hamster kidney (BHK21) cells, human HeLa cells, human retinal pigment epithelial (RPE-1) cells and rat embryo fibroblasts (REF52) were cultured as described elsewhere (Marie et al. 2012; Palokangas et al. 1998).

### Transferrin uptake

To provide fluorescent labeling of the endocytic membrane recycling compartments via uptake of transferrin the cells were washed twice with prewarmed 37°C DMEM supplemented with 0.2% bovine serum albumin (BSA) and 10 mM HEPES, pH 7.2, followed by incubation for 60 min at 37°C in the same serum-free medium. For transferrin uptake, cells were incubated for an additional 60 min in the serum-free medium containing 20 μg/ml of human transferrin coupled to Alexa Fluor 488 or Alexa Fluor 594.

### Experimental treatments

To release membrane-bound COPI coats and induce complete breakdown of the Golgi apparatus, NRK cells were incubated for 30 min at 37°C in medium containing 5 μg/ml BFA. The disassembly of MTs and the spindle apparatus was obtained by 30 min treatment with 10 μg/ml nocodazole (NZ). To selectively disassemble biomolecular condensates, the cells were incubated for 1, 2.5 or 5 min at 37°C in culture medium supplemented with 2.5% propylene glycol (PG). Upon wash-out the cells were washed twice with 37°C culture medium, followed by incubation for an additional 30 min in the absence of PG. When the effect of low temperature on the mitotic condensates was examined, the coverslips were immersed for 1, 2.5 or 5 min in ice-cold culture medium prior to fixation.

### Fixation conditions

In the standard sample preparation protocol the cells grown on glass coverslips were fixed for 60 min with 3% paraformaldehyde (PFA) in 0.1 M Na-phosphate buffer (pH 7.2) at RT. To obtain better structural preservation of the mitotic condensates the cells were fixed for 120 min with ice-cold PFA-lysine-sodium periodate (PLP) fixative (McLean and Nakane 1974), consisting of 2% PFA, 0.075 M lysine-HCl and 0.01 M NaIO_4_ in 0.375 M Na-phosphate buffer, pH 6.2). This fixative preserves cell morphology better by cross-linking carbohydrates, retaining at the same time the antigenicity of proteins (Brown and Farquhar 1989). The coverslips were quickly immersed in ice-cold fixative and kept on ice for the first 30 min, followed by transfer to RT for the remaining 90 min. In some experiments cells were further fixed and permeabilized by incubation for 5 min at -20°C in ice-cold methanol.

### Immunofluorescence staining and confocal microscopy

In most experiments, the fixed cells were permeabilised using 0.2% saponin with saponin being present throughout the labelling protocol. The immunofluorescence staining protocol, including the exposure of antigenic sites using guanidine-HCl (Peränen, Rikkonen, and Kääriäinen 1993) has been described in detail previously (Raddum et al. 2013; Sannerud et al. 2008). After staining, the cells were first examined in a Zeiss Axiovert 200M inverted microscope equipped with long-working distance Plan-NEOFLUAR 40X and 100X objectives and fluorescence filter appropriate for Alexa 488, Alexa 594 and DAPI. Confocal microscopy on selected specimens was carried out to obtain individual optical sections or Z-stacks (step size 0.3 or 0.5 μm) using a Leica SP5 AOBS or Leica TCS SP8 confocal laser scanning microscopes, equipped with a 63X/1.4 NA Plan-Apochromat or 100X NA1.4 HC PL APO STED White oil-immersion objectives, _∼_1 Airy unit pinhole aperture, 405 Diode, Argon and Helium/ Neon lasers and the appropriate filter combinations. The images prepared using ImageJ are presented as single sections or maximum-intensity projections. Imaris software was used for image processing and preparation of the animations (**Supplementary movies 1 and 2**).

### Quantitations

The Zeiss Axiovert 200M fluorescence microscope equipped with the 100X objective was used to determine the percentages of cells at different phases of mitosis or cytokinesis, positive for the large AnxA2-positive structures, by examining a total of ca. 600 mitotic cells (**Fig. 1B**). The same microscope was employed in the investigation of the effect of the aliphatic alcohol PG on the mitotic structures, based on the analysis of 50–85 meta- or anaphase cells for each test point (**Fig. 4A**). To establish the average size of the mitotic condensates in prometa-, meta- and anaphase cells (**Fig. 1C**), Z-stacks were generated by the Leica TCS SP8 confocal microscope, followed by measurement of the diameters of up to 100 structures for each mitotic stage from maximum intensity projections corresponding to the center of the cell. The numbers of cells subjected to confocal optical sectioning to calculate the approximate number of condensates in cells at different stages of mitosis (**Fig. 6A**) were as follows: prometaphase (n=4), metaphase (n=7), early anaphase (n=5) and late anaphase (n=13).

## Acknowledgements

We are grateful to Dr. Volker Gerke and Dr. Kristian Prydz for critical reading and constructive comments on the manuscript. This study was supported by the University of Bergen, The Western Norway Regional Health Authority (grant no. 911499 to AV), The Research Council of Norway (grant no. 240400/F20 to AV and grant 196745/V45 to JS and Kristian Prydz) and the Fridtjof Nansen Foundation (JS). The funding sources were not involved in the design of the experiments, writing of the paper, or presentation of the data. The microscopy part of the work was carried out at the Molecular Imaging Centre (MIC) at the Department of Biomedicine, University of Bergen.

## Conflict of interest

The authors declare that they have no conflict of interest.

## Author Contributions

Conceived and designed the experiments: AKG, JS and AV. Performed the experiments: AKG, HD, JN, SSP and JS. Analyzed the data: AKG, JS and AV. Wrote the paper: AKG, AV and JS

## Abbreviations

AnxA2: Annexin A2 protein;
BFA: brefeldin A;
BHK21: baby hamster kidney cells;
COPI: coat protein I;
DAPI: 4’,6-diamidino-2-phenylindole;
EEA1: early endosomal antigen 1;
ER: endoplasmic reticulum;
HeLa: human cervical cancer cells;
IC: intermediate compartment;
LLPS: liquid-liquid phase separation;
MTs: microtubules;
NE: nuclear envelope;
NRK: normal rat kidney cells;
NZ: nocodazole;
PC12: rat pheochromocytoma cells;
PCM: pericentriolar material;
PI(4,5)P_2_: phosphatidyl-inositol-4,5-bisphosphate;
PFA: paraformaldehyde;
PG: propylene glycol;
PLP: periodate-lysine-paraformaldehyde;
PM: plasma membrane;
PML: promyelocytic leukemia;
REF52: rat embryonal fibroblasts;
REs: recycling endosomes.

## Supplementary figure legends

**Figure S1.** *Human HeLa cells lack the AnxA2-containing mitotic structures*. HeLa cells were subjected to endocytic uptake of Alexa-Fluor 594-coupled Tfn (red), fixed with PLP and stained with antibodies against AnxA2 (green) and DAPI. Note the diffuse localization of AnxA2 at the cell cortex, the presence of endocytosed Tfn in peripheral structures of variable size (white arrows) and its piling-up at the spindle poles of the metaphase cell (opens arrows). Scale bar: 10 µm.

**Figure S2.** *The mitotic condensates do not contain ER or Golgi membranes*. NRK cells expressing GFP-Rab1 were fixed with PLP and stained with antibodies against calnexin (ER; panels A-C), GM130 (*cis*-Golgi; panels D-F), mannosidase II (*medial*-Golgi; panels G-I) or TGN38 (*trans-*Golgi/TGN; panels J-L). Note the absence of all these secretory compartment markers from the Rab1-positive peripheral structures of metaphase cells. The diffuse staining for mannosidase II corresponds to the vesicular “Golgi haze”, while both GM130 and TGN38 partially colocalize with the Rab1-positive IC elements at the spindle poles (arrows). An interface cell is marked with an asterisk in panels A-C. Scale bars: 10 µm.

**Figure S3.** *The mitotic structures are not affected by brefeldin A (BFA).* Control NRK cells expressing GFP-Rab1 (panels A-C), or cells treated for 30 min with BFA (5 µg/ml) (panels D-F) were fixed with PLP and stained for AnxA2 (red). BFA does not appear to influence the structure or localization of the large peripheral structures, or appearance of the condensed chromosomes in metaphase cells, but by releasing membrane-bound COPI coats results in the tubulation of the IC elements at the spindle poles (open arrow). Scale bars: 10 µm.

**Figure S4.** *The mitotic structures contain Rab11-positive REs but lack early or late endosomes or lysosomes.* The localizations of different endosomal markers, including Rab7 (panels A-C), Rab11 (panels D-F), as well as EE1 and LAMP-1 (panels G*-*I and J-L) in cells at metaphase were compared with markers of the large peripheral structures – transferrin (Tfn), lamin B and Rab1, respectively. Note that in contrast to the RE marker Rab11, the early (EEA1) or late endosomal/lysosomal markers (Rab7, LAMP-1) cannot be localized to the large mitotic structures. Interphase cells are marked with asterisks. Scale bars: 10 µm.

## Supplementary movies

**Movie 1.** *Animation demonstrating the peripheral localization of the mitotic condensates in a metaphase cell*. NRK cells expressing GRP-Rab1 were stained with antibodies against AnxA2 and with DAPI. The Rab1- and AnxA2-positive mitotic structures (yellow) and condensed chromosomes (blue) were surface-rendered, and the movie was prepared using the Imaris software. Note also the localization of the Rab1-containing IC elements (green) at the spindle poles.

**Movie 2.** Movie 1. *Animation showing mitotic partitioning of the condensates.* NRK cells expressing GRP-Rab1 were stained with antibodies against AnxA2 and with DAPI. The condensed chromosomes (blue) and the Rab1- and AnxA2-positive large mitotic structures in the forming daughter cells were surface-rendered, and the latter were differentially pseudo-coloured. The movie was prepared using the Imaris software.

## Notes

### Competing Interest Statement

The authors have declared no competing interest.

