## Supplemental Figures 1-4 for "Annexin A2 and lamin B join membrane recycling compartments for the assembly of biomolecular condensates operating in mitotic partitioning"

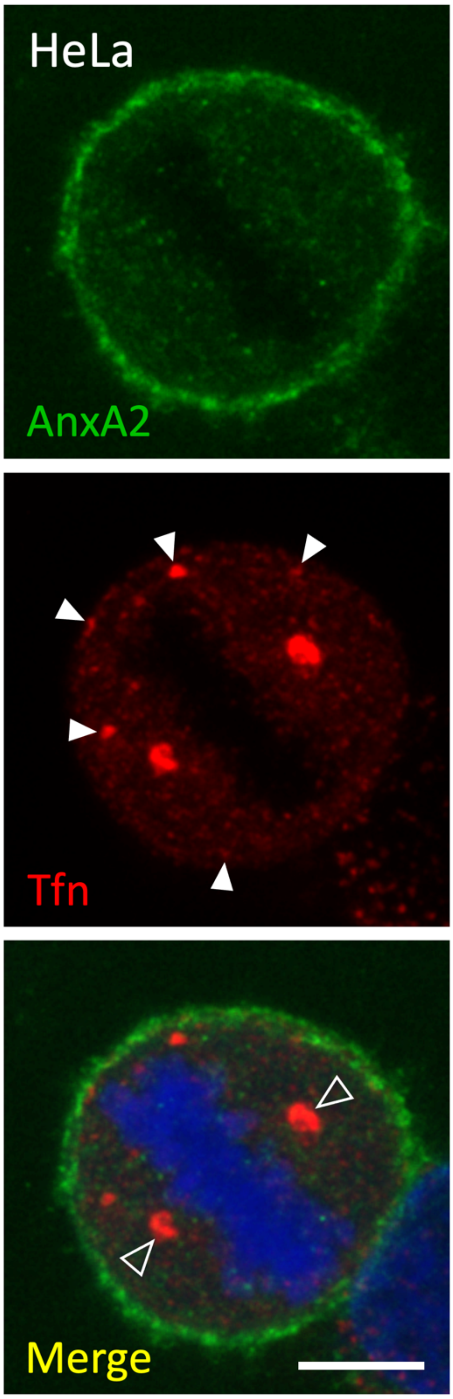


**Supplementary Figure S1**

Grindheim et al.

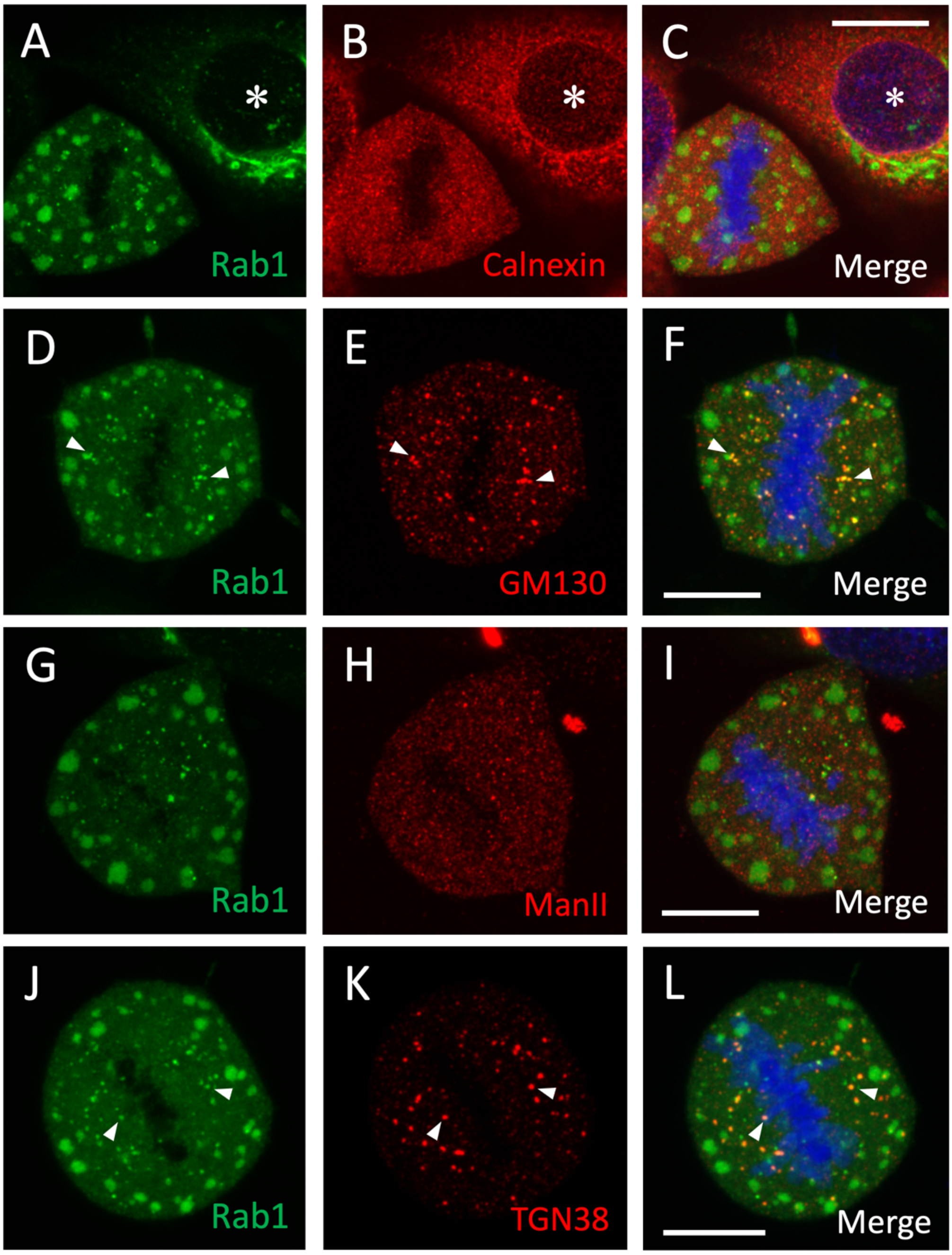


**Supplementary Figure S2**

Grindheim et al.

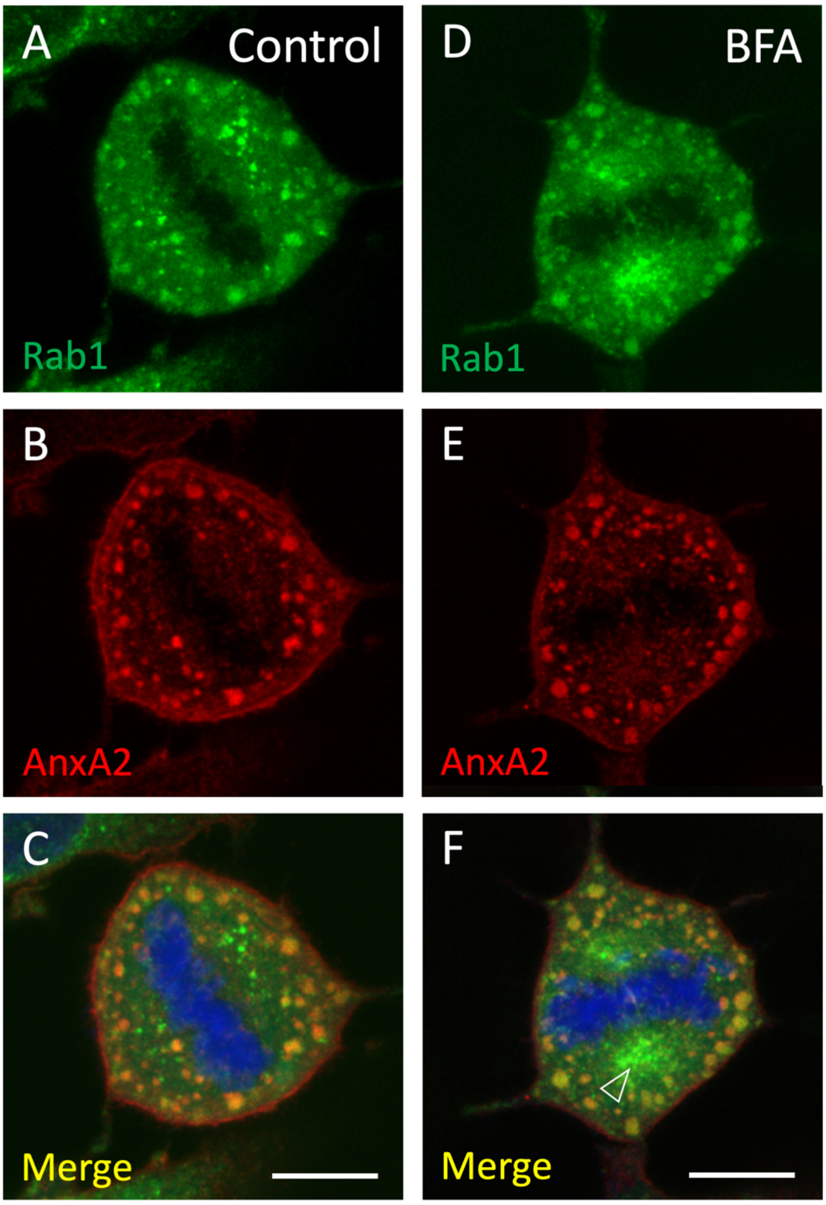


**Supplementary Figure S3**

Grindheim et al.

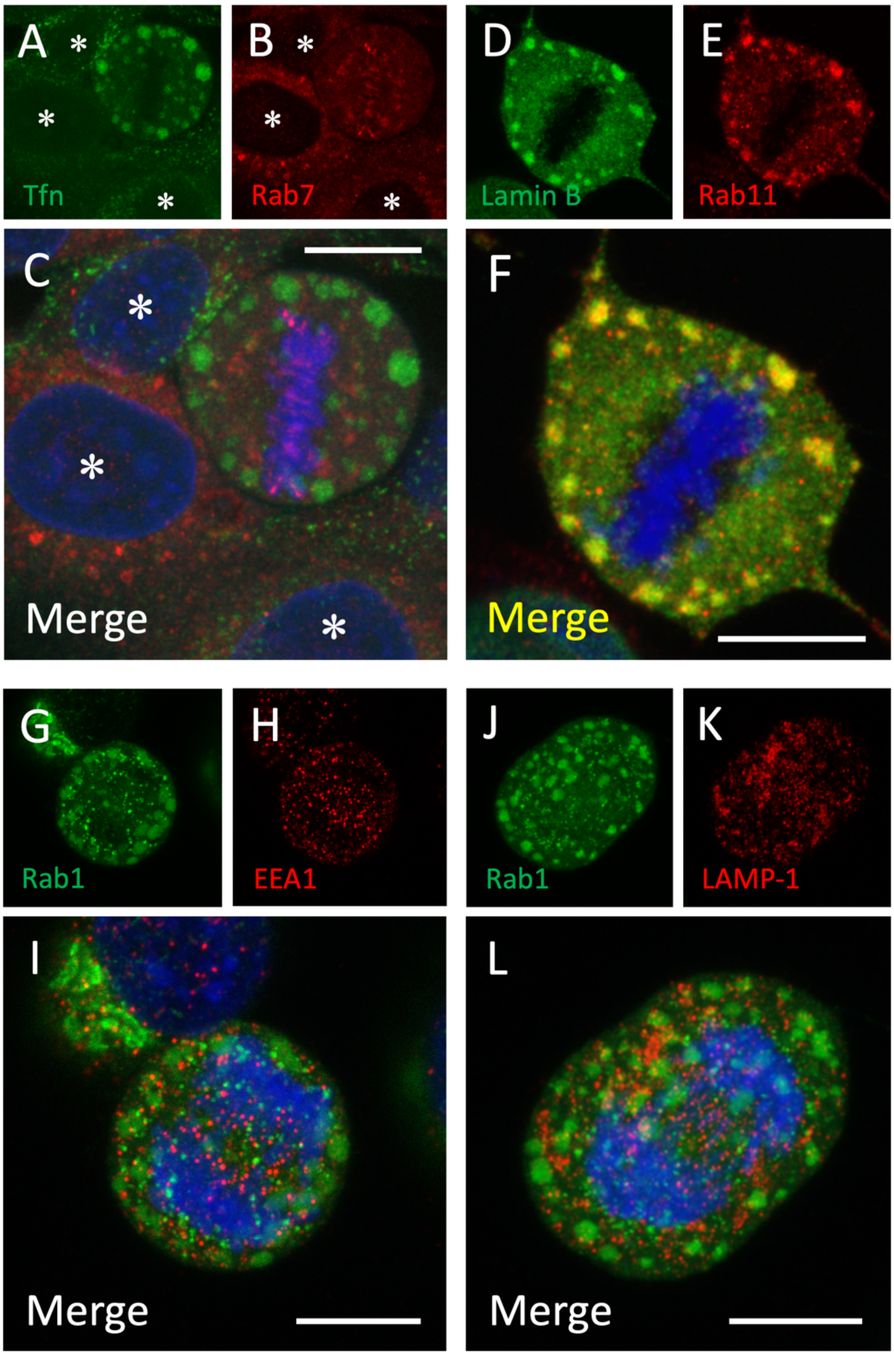


**Supplementary Figure S4**

Grindheim et al.
